# Detecting differential alternative splicing events in scRNA-seq with or without UMIs

**DOI:** 10.1101/738997

**Authors:** Yu Hu, Kai Wang, Mingyao Li

## Abstract

Analysis of alternative splicing in single-cell RNA sequencing (scRNA-seq) is challenging due to its inherent technical noise and generally low sequencing depth. We present SCATS (Single-Cell Analysis of Transcript Splicing) for differential alternative splicing (DAS) analysis for scRNA-seq data with or without unique molecular identifiers (UMIs). By modeling technical noise and grouping exons that originate from the same isoform(s), SCATS achieves high sensitivity to detect DAS events compared to Census, DEXSeq and MISO, and these events were confirmed by qRT-PCR experiment.

---

The emergence of scRNA-seq technology has made it possible to measure gene expression variations at cellular level^1^. This breakthrough enables the investigation of a wide range of problems including analysis of splicing heterogeneity among individual cells. However, compared to bulk RNA-seq, scRNA-seq data are much noisier due to high technical variability, low sequencing depth, and the lack of full length transcript sequencing for droplet based protocols. Despite the growing popularity of scRNA-seq, few published studies have investigated alternative splicing, and even when studied, methods developed for bulk RNA-seq were utilized^2, 3^, which may not be optimal for scRNA-seq data.

Methods designed specifically for splicing analysis in non-UMI based scRNA-seq data only started to emerge recently^4^. Huang *et al*.^5^ detects differential exon-usage by performing a pairwise comparison between every two cells, which becomes computationally infeasible when large number of cells are generated from droplet-based protocols^6-8^. Song *et al.*^9^ quantifies exon-inclusion levels based on junction-spanning reads exclusively. However, these estimations are unreliable due to sparse read counts that span exon-exon junctions and technical noise of scRNA-seq data, leading to limited power in detecting DAS events. Qiu *et al.*^10^ and Ntranos *et al.*^11^ developed approaches to detect differential transcript usage based on pre-estimated cell-specific isoform expressions or transcript compatibility counts. Although encouraging, the feasibility of estimating isoform usage at single-cell level still remains questionable due to limited informative reads for splicing in scRNA-seq.

Here we present SCATS, which achieves high sensitivity to detect DAS events in scRNA-seq by accounting for technical noise and low sequencing depth, through an exon-grouping approach originally developed in PennDiff^12^. Given annotated transcript information, sequencing reads aligned to exons originated from the same isoform(s) are first grouped together (**Fig. 1a**). This grouping step is essential in splicing analysis of scRNA-seq data as it naturally aggregates spliced reads across different exons, making it possible to detect DAS events even for genes with low sequencing depth^12^. Moreover, SCATS can be applied to scRNA-seq data either with or without UMIs. For non-UMI data, SCATS explicitly models technical noise by accounting for capture efficiency, amplification bias and drop out events through the use of external spike-ins; for UMI data, SCATS models capture efficiency and further accounts for transcriptional burstiness (**Fig. 1b-d**) in detecting DAS events between groups of cells.

**Fig. 1.**
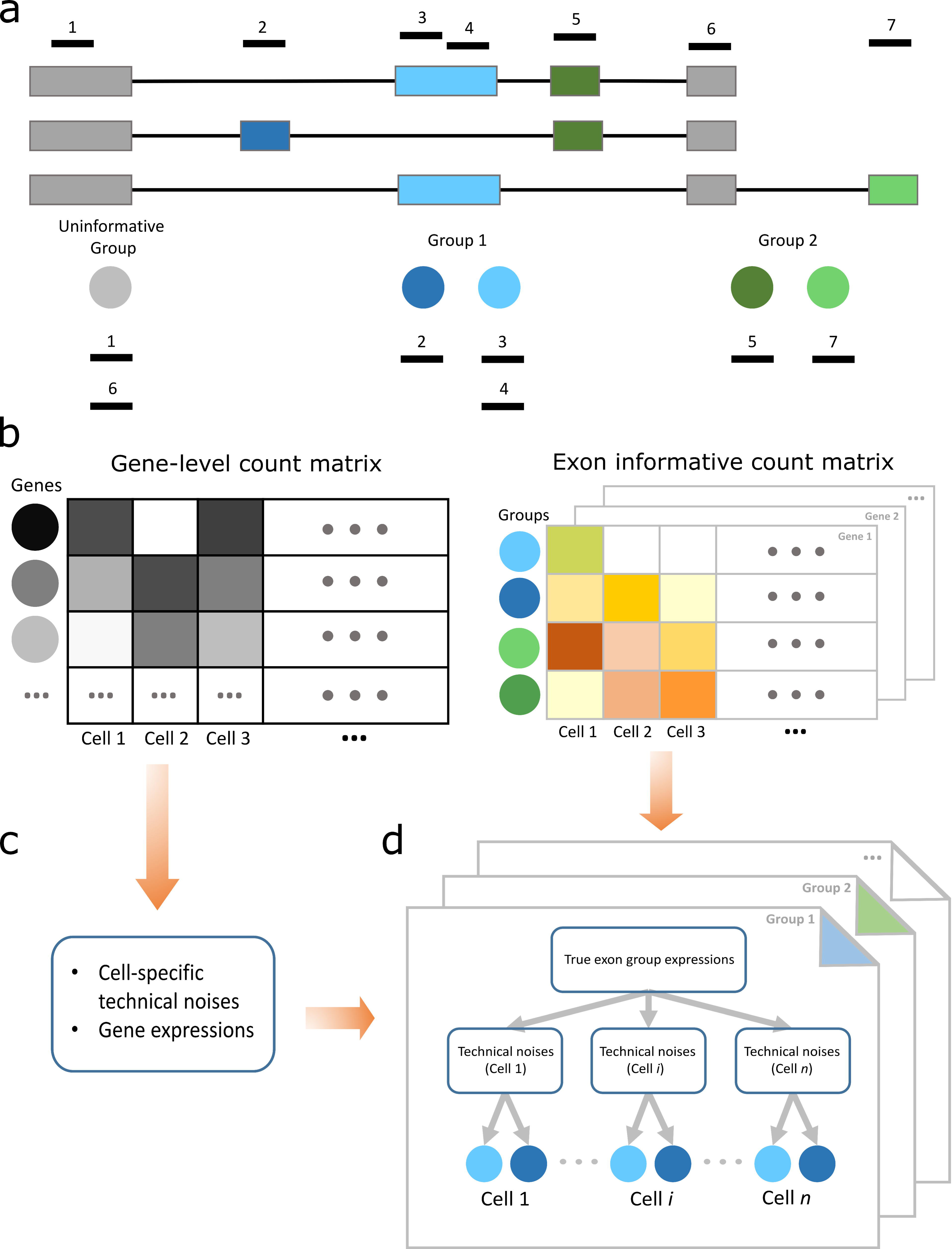
Workflow of SCATS. **(a)** Exons were first divided into groups based on their isoform origin. Those that originate from the same isoform(s) are grouped together and share the same exon-inclusion level. Group 1 (blue) and group 2 (green) are informative for alternative splicing, while uninformative group (grey) is not. Within each informative group, exons were further classified to included (inclusion level = *ψ)* and excluded (inclusion level = *ψ*) region. Dark blue and dark green represent included regions. Light blue and light green represent excluded regions. **(b)** Observed read counts of scRNA-seq. Gene-level read count matrix (left) and group-level informative read count matrices (right) are summarized from aligned scRNA-seq data in BAM format. **(c)** Given gene-level read count matrix, cell-specific technical noises and population level gene expressions are quantified using TASC^18^. **(d)** Given technical parameter estimates from **(c)**, for each alternative splicing exon group, the statistical inference (exon-inclusion level estimation and inclusion level difference testing) on exon-inclusion levels was made based on a hierarchical model that accounts for technical noise.

Exon-inclusion level, which measures the relative usage of an exon, is a commonly used measure to quantify the process of alternative splicing. A critical step in DAS detection is to reliably estimate exon-inclusion levels for exons that originate from the same isoform(s). Our simulations indicate that SCATS’s estimates are strongly correlated with the true levels (R = 0.83), whereas naïve estimation that ignores technical noise in scRNA-seq only leads to a correlation of 0.66 (**Supplementary Fig. 2a**). We further assessed the performance of SCATS in detecting DAS events, and compared with two other state-of-the-art methods: Census^10^, an algorithm designed to preprocess raw read counts by removing technical noise, and DEXSeq^13^, a popular method for bulk RNA-seq splicing analysis. Our results indicate that SCATS has well controlled type I error rates (**Supplementary Fig. 2b,c**), whereas Census tends to yield false positive results with enrichment of p-values near zero, and DEXSeq is overly conservative with enrichment of p-values near one. SCATS is also more powerful than Census and DEXSeq (**Supplementary Fig. 3**), although Census has severely inflated type I error rates. Compared to DEXSeq, SCATS performs consistently better, especially when the true exon-inclusion level difference is small. This indicates that SCATS is robust in detecting subtle splicing difference, thanks to the accurate exon inclusion-level estimation due to exon grouping and technical noise modeling.

Next, we applied SCATS to a scRNA-seq dataset generated from adult mouse brains, which includes 1,679 cells generated using the SMART-seq protocol (Tasic *et al.* dataset^14^) (see details in **Methods**). This dataset includes three major cell classes and 49 sub cell types (23 GABAergic cell types, 19 glutamatergic cell types, and seven non-neuronal cell types). Major cell classes and sub cell types reported in the original publication were treated as ground truth to evaluate SCATS in characterizing splicing heterogeneity across cells. This dataset is non-UMI based, but read counts from 92 ERCC spike-ins are available, which allow us to quantify technical variations. To assess the accuracy of exon-inclusion level estimation of SCATS, we selected two consecutive alternatively spliced exons (flip-flop exons^15^) from two Glutamate receptor genes *Gria1* and *Gria2* (**Supplementary Fig. 4a,b**), which have been reported to display a highly variable splicing pattern across cell types^14, 15^. These flip-flop exons are complementary with each other, and each single one is included in a different mature RNA. Sommer *et al.*^15^ measured the expression levels of the flip and flop exons, respectively, with RNA *in situ* hybridization (ISH). Their findings showed that the flip exon was preferentially utilized in layer II/III of neocortex as compared to the flop exon, whereas both exons were equally expressed in layer VI of neocortex. To evaluate the exon usage quantification of SCATS, we estimated cell type-specific exon-inclusion level of the flip exon. As shown in **Fig. 2a,b**, the estimated exon-inclusion levels of the flip exon in cell types L2-Ngb (layer II) and L2/3-Ptgs2 (layer II / III) are over 80%, whereas the estimates are around 50% in cell type L6a (layer VI) (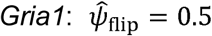 and 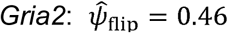). These results are consistent with RNA ISH measurements in Sommer *et al*.^15^.

**Fig. 2.**
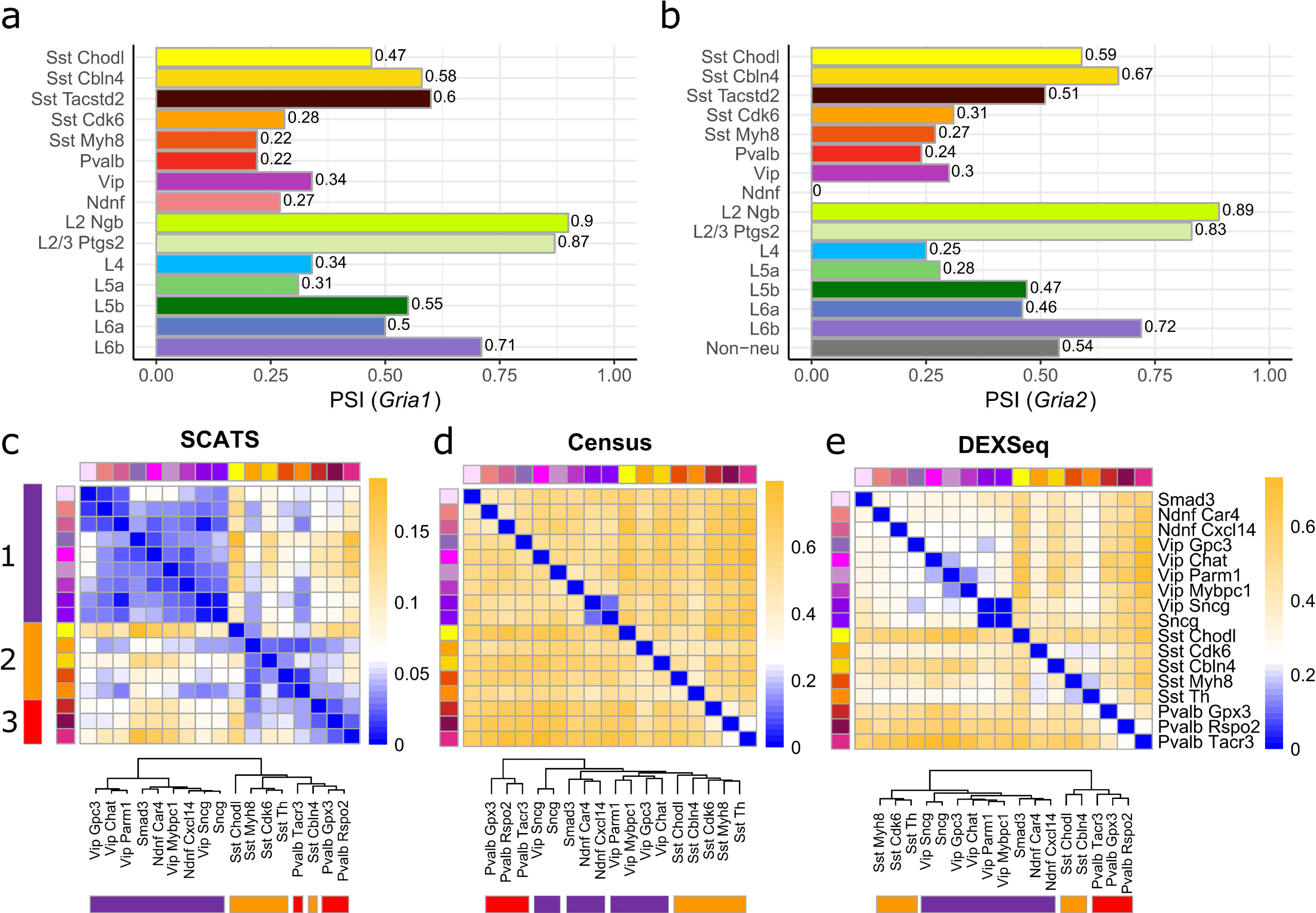
DAS analysis results of the Tasic *et al.* data. **(a,b)** Alternative splicing exon quantification of flip and flop exons from gene *Gria1* and *Gria2* using SCATS. Barplots showing exon-inclusion level estimates of the flip exon from gene *Gria1* **(a)** and *Gria2* **(b)** across different cell types. Colors indicate different cell types. These estimates agree well low-resolution RNA ISH data. **(c-e)** Pairwise DAS comparison across GABAergic cell types from mouse cortex using SCATS **(c)**, Census **(d)** and DEXSeq **(e)**. Colors indicate different GABAergic cell types. Purple, yellow and red indicate three sub cell classes. Heatmap showing the proportion of detected DAS exons groups for each pairwise comparison between cell types based on 966 exon groups from 296 marker genes. Dendrogram depicting cell classification results of the 17 GABAergic cell types. The distance metric between two cell types is the proportion of detected DAS exons by corresponding approach: SCATS **(a)**, Census **(b)** or DEXSeq **(c)**. SCATS outperforms in identifying splicing heterogeneity across different cell types, while controlling false positive rate.

Encouraged by the accurate exon-inclusion estimates, we next conducted a pairwise DAS analysis across all 49 sub-cell types based on 296 most differentially spliced genes (with 966 exon groups) selected by Tasic *et al*. using MISO^16^, a method developed for splicing analysis in bulk RNA-seq data. **Supplementary Fig. 5** shows the quantile-quantile (QQ) plots of the log-transformed p-values from pairwise comparisons of different sub cell types, which include sub-cell type comparisons both within major cell classes and cross major cell classes. For tests within GABAergic, Glutamatergic and non-neurons, the p-values from SCATS are distributed reaches to slightly below the red diagonal line until when the significance level *α* reaches to 0.01. This pattern is consistent with the QQ plot shown in the simulations **(Supplementary Fig. 2b)**. In cell class subsets, indicating that more DAS exons were detected at less stringent significance contrast, there is a sharp deviation from the diagonal line at *α =* 10^−15^ for all three cross-major thresholds. This is not surprising because more DAS exons are expected to be detected between sub-cell types from different major cell classes compared to cell types that originate from the same major cell class. **Supplementary Fig. 6a,b** show DAS heterogeneity revealed by SCATS across the 49 sub cell types based on the 966 exon groups in these 296 genes. The heatmap shows three clusters, consistent with the three labeled major cell classes: GABAergic, glutamatergic and non-neuronal. Although this pattern was also revealed by MISO analysis in Tasic *et al*., SCATS results revealed more heterogeneous splicing patterns within each major cell class than MISO, demonstrating its higher sensitivity in detecting splicing heterogeneity.

For GABAergic cells, Tasic *et al.* conducted a hierarchical clustering analysis based on qRT-PCR measurements obtained from 79 marker genes for GABAergic sub cell types. These cells were classified into three sub-clusters. To further assess the accuracy of SCATS in profiling splicing heterogeneity, we treated these qRT-PCR based clustering result as ground truth and asked whether SCATS can reveal this sub-clustering structure from its detected DAS events. MISO results in Tasic *et al.* showed homogenous DAS pattern across GABAergic cells with no cell subpopulations being clustered together. Heatmaps in **Fig. 2c-e** show DAS analysis results of the 480 GABAergic cells using SCATS, Census and DEXSeq, respectively. The three sub-clusters (purple, orange, red) were identified by SCATS but missed by Census and DEXSeq. These results demonstrate the high sensitivity of SCATS in detecting DAS among closely related cells from the same major cell class. Additionally, we generated a sashimi plot using IGV to validate the detected DAS exons by SCATS. **Supplementary Fig. 9** shows the empirical evidence of a DAS event at the flip-flop exons between Sst-Cdk6 and Sst-Cbln4 for gene *Gria1*, which is consistent with SCATS (*P* = 0.018).

Although designed primarily to detect DAS events in full-length transcript scRNA-seq data, SCATS can also detect DAS events for UMI data, which typically only contain sequences from the 3’ or 5’ end of a transcript. To evaluate the performance of SCATS when full-length transcript information is not available, we analyzed the Zeisel *et al*. dataset^17^, which are UMI-based data generated from 3,005 cells in mouse cortex. This dataset includes nine major cell classes and 47 sub cell types. We performed DAS analysis on 1,826 marker genes (3,542 exon groups) for these nine major cell classes across the 47 sub cell types in this dataset. Since UMI counts were consolidated from non-UMI reads, we compared read count distributions between UMI and non-UMI data. Not surprisingly, splicing informative read counts in UMI data are much sparser than non-UMI data (**Supplementary Fig. 10a,b**), making it challenging to detect DAS in UMI data. However, by exon grouping and appropriate statistical modeling, SCATS is able to detect DAS even with such sparse data. **Fig. 3a** shows the heatmap of the proportions of the detected DAS events for each pairwise comparison by SCATS. Neuronal cells are clearly differentiated from non-neuronal cells, although some major cell classes (marked by single colors) cannot be discerned from this heatmap. To better understand these results, we treated the proportions of significant DAS exons for each pairwise comparison as a similarity metric and performed hierarchical clustering analysis as shown in **Fig. 3b**. For SCATS, the separation in neuron cells agrees well with labeled cell types, with only one CA1 Pyramidal cell type (ClauPyr) was misclassified as Interneurons. For non-neuronal cells, SCATS clearly separated the Oligodendrocytes and Astrocytes cell classes from others, with four labels in Ependyma, Microglia and Mural misclassified. These clustering results demonstrate the reliability of DAS detections across cell types using SCATS even when informative read counts are sparse. We also conducted similar pairwise DAS analyses using MISO, Census and DEXSeq. Census may yield more false positive results because it detected more than 40% DAS events across most pairwise comparisons, even for those comparisons within major cell classes (**Fig. 3c,d**). In contrast, DEXSeq and MISO missed most of the DAS events and the hierarchical clustering analysis failed to separate different major cell types for both neurons and non-neurons (**Fig. 3e,f, Supplementary Fig. 10c,d**). This is not surprising as informative molecule counts in UMI-based scRNA-seq data are extremely sparse, and methods developed for bulk RNA-seq data are not optimal when analyzing scRNA-seq data. It is worth noting that splicing quantification offers higher resolution of cellular heterogeneity than total gene expression. As shown in **Supplementary Fig. 10e**, exon-inclusion levels showed more distinct patterns across cell types for gene *Gria1* than total gene expression.

**Fig. 3.**
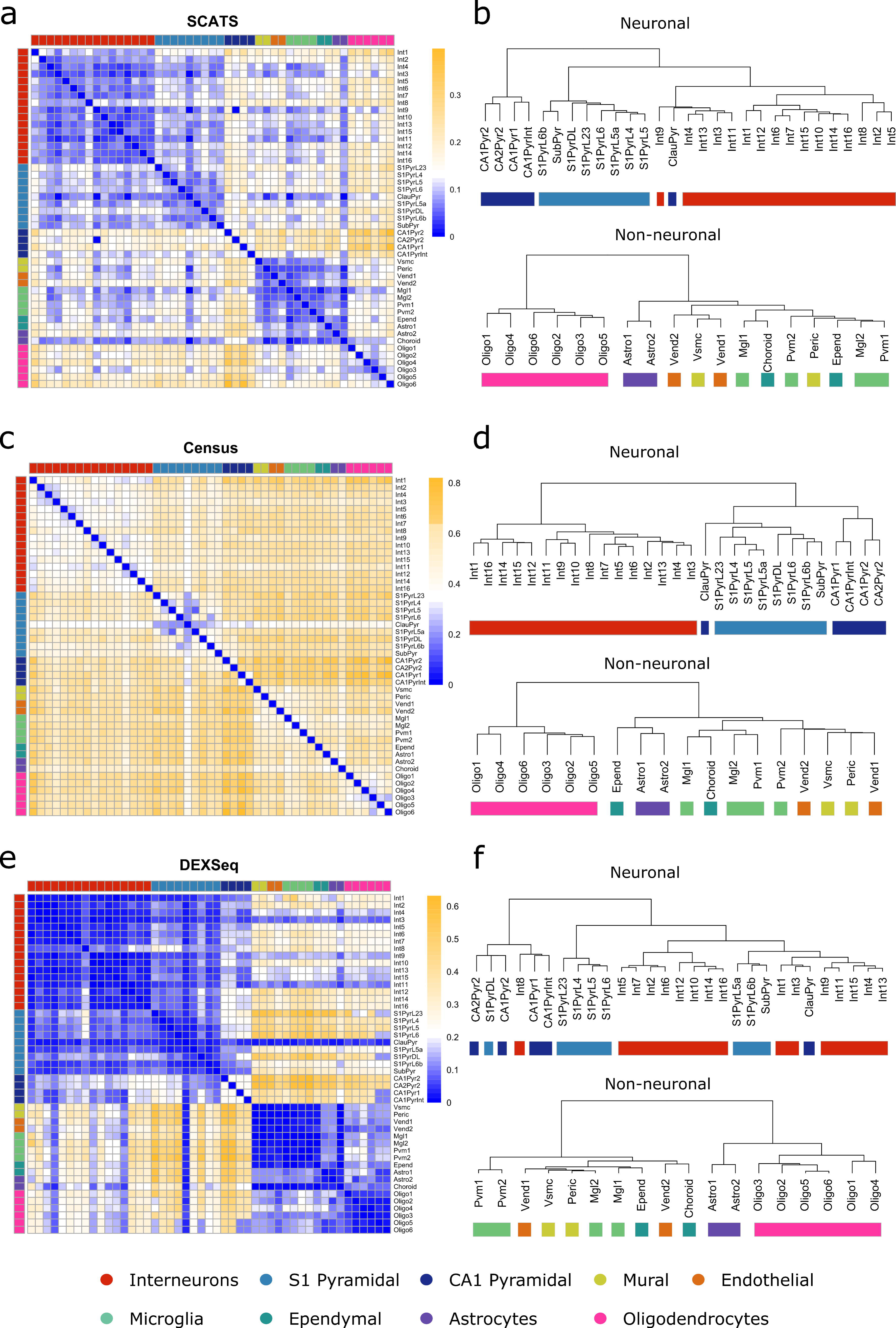
DAS analysis results of the Zeisel *et al*. data. **(a-f)** Pairwise DAS comparison across 3,005 cells (nine major cell types) from mouse cortex and hippocampus using SCATS **(a,b)**, Census **(c,d)** and DEXSeq **(e,f)**. Colors indicate nine major cell types. **(a,c,e)** Heat map showing the proportion of detected DAS exons groups (3,542 exon groups from 1,826 genes) for each pairwise comparison between cell types. **(b,d,f)** Dendrogram depicting cell classification results of the 47 sub cell types. The distance metric between two cell types is the proportion of detected DAS exons among all analyzed exons by each approach: SCATS **(b)**, Census **(d)** or DEXSeq **(f)**. Zeisel *et al.* data shows highly cell type-specific alternative splicing pattern.

ScRNA-seq data offer new insights into alternative splicing at cellular level, which helps us to understand cellular heterogeneity with higher resolution as compared to bulk RNA-seq data. However, low sequencing depth and technical noise have precluded the investigation of splicing heterogeneity in most scRNA-seq studies. To fill in such knowledge gap, we developed SCATS, which utilizes informative read counts at exon level to achieve high sensitivity in DAS detection. By grouping exons that originate from the same isoform(s) and modeling of technical noise, SCATS aggregates spliced reads across different exons, making it possible to detect splicing events even when sequencing depth is low. With the increasing adoption of scRNA-seq, we believe SCATS will be well-suited for various splicing studies and offers additional insights beyond total gene expression analysis.

## ACKNOWLEDGEMENTS

This work was supported by the following grants: NIH R01GM108600, R01GM125301, R01HL113147, and R01EY0301092 (to M.L.).

## AUTHOR CONTRIBUTIONS

This study was conceived of and led by M.L.. H.Y. and M.L. designed the model and algorithm. H.Y. implemented the SCATS software and led the data analysis with input from M.L. and K.W.. H.Y. and M.L. wrote the paper with feedback from K.W..

## COMPETING FINANCIAL INTERESETS STATEMENT

The authors declare no competing interests.

## METHODS

### Splicing informative read counts based on exon grouping

To get informative reads for DAS analysis, we pre-process sorted isoform annotation file in GTF format and sorted aligned scRNA-seq data in SAM format. For each gene, alternatively spliced exons are grouped if they originate from the same isoform(s). Suppose the gene under consideration has *M* exon groups, then all reads mapped to this gene can be grouped into 2*M*+1 categories: 1) uninformative reads: reads that map to exons shared by all isoforms of the gene; 2) included (+) informative reads of exon group *j*: reads mapped to alternatively spliced (AS) exons that are included in exon group *j* (1 ≤ *j* ≤ *M*); and 3) excluded (-) informative reads of exon group *j*: reads mapped to AS exons that are excluded from exon group *j* (1 ≤ *j* ≤ *M*).

Let ***Y***_*i*_ = (*Y*_*i*0_, *Y*_+*i*1,…_ *Y*_+*iM*,…_, *Y*_−*i*1,…,_ *Y*_−*iM*_) be the observed read counts in cell *i* for the gene, where *Y*_*i*0_ denotes uninformative read count, *Y*_+*ij*_ denotes included informative read counts of exon group *j*, and *Y*_−*ij*_ denotes excluded informative read counts of exon group *j*. **Fig. 1a** shows an example of how exons are grouped and informative reads are counted. For this hypothetical gene, there are three isoforms constructed by five exons, where three of them are alternatively spliced and grouped into two groups (blue and green). Among the seven mapped reads, reads 1 and 6 are uninformative reads (*Y*_+*i*0_ = 2), read 2 is an informative read included in exon group 1(*Y*_+*i*1_ = 1), reads 3 and 4 are informative reads excluded from group 1(*Y*_+*i*0_ = 2), read 5 is an informative read included in exon group 2(*Y*_+*i*2_ = 1), and read 7 is an informative read excluded from group 2(*Y*_−*i*2_ = 1).

### Modeling splicing informative UMI and non-UMI counts

SCATS is designed to detect DAS events at exon group level. For the ease of notation, we drop the exon group index. For a given exon group of interest, let *ψ* be the inclusion level of an included (+) exon group and *θg* be the mean expression of gene *g*. To account for cell-specific technical noise, we model included informative read counts *Y*_+*i*_ and *Y*_−*i*_ using a hierarchical model (**Supplementary Fig. 1**). We assume cell-specific true expression level of the included/excluded exon group *μ*_+*i*_ /*μ* _−*i*_ follows a log-normal distribution:

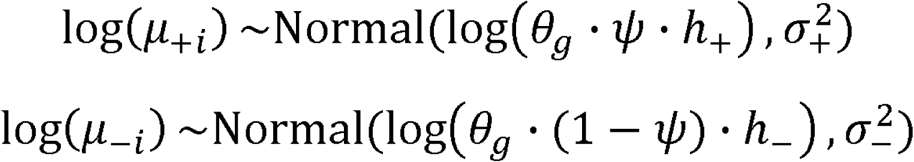

where *h*_+_/*h*_−_ represents the probability of included/excluded reads being informative and 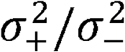 is variance of expression level *μ*_+*i*_ /*μ* _−*i*_ in log scale.

For non-UMI count, amplification bias and low capture efficiency in scRNA-seq are quantified by a linear model. Specifically, for cell *i*, the expected value of the informative read count *λ* _+*i*_/*λ*_−*i*_ is modeled as

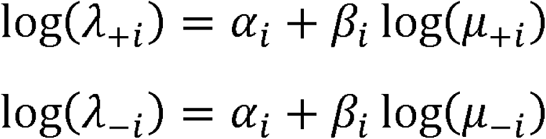

where *α*_*i*_ and *β*_*i*_ are cell-specific technical parameters designed to model capture efficiency and amplification bias, respectively. These technical parameters can be quantified with TASC^18^ when ERCC spike-ins are provided.

For UMI count, cell-specific parameters *β*_*i*_ are equal to 1 as amplification bias can be alleviated by UMI. Therefore, the expected informative UMI count *λ* _+*i*_/*λ*_−*i*_ is proportional to true expression of the included/excluded exon group:

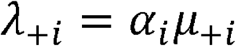

Moreover, we account for inflated zeros of scRNA-seq which are primarily sourced from two contributing factors: technical drop-out and transcriptional bursting. Drop-out happens when a particular transcript is lost in scRNA-seq experiment steps while the gene is expressed in the cell. Also, excessive zero counts can be introduced when significant bursting genes are in “off” state due to biological heterogeneity. Previous studies have shown that the modeling of drop-out in UMI data is unnecessary^19, 20^. Therefore, to model zero inflation, for a given gene, we let *Z*_*i*_ be the indicator that the gene in cell *i* is captured in the library (drop-out does not happen) and it is in the “on” state. The probability of *Z*_*i*_ = 1 is *π*_*i*_. Therefore, given *λ* _+*i*_ and *λ*_−*i*_, the distribution of the observed informative read count *Y*_+*i*_/*Y*_−*i*_ is

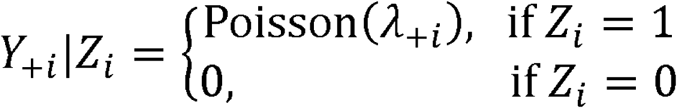

And

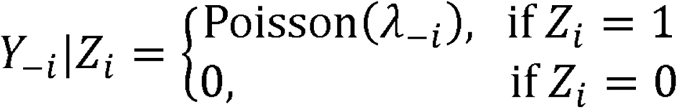

where Poisson distribution has been reported to approximate the read sample process well after zero inflation being removed^19, 20^.

### Probability of included/excluded reads being informative

Another key part of SCATS is to estimate *h*_+_/*h*_−_, the probability of included/excluded reads being informative. From the isoform structure of gene *g*, we can determine the origin (exon group) of a read (being informative read) only when it is mapped to exon-exon junction or an alternatively spliced exon. Thus, a natural way to estimate the probability that a read being mapped to this region is to calculate the ratio between weighted length of the informative region and the whole region. Due to low sequencing depth of scRNA-seq data, we assume the exon group shares the same *h*_+_/*h*_−_ across cells instead of estimating cell-specific parameters:

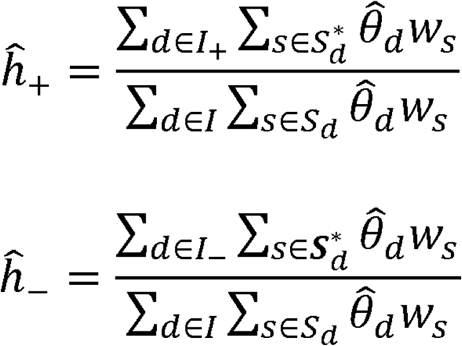

where *I*_+_ and *I*_−_denote the set of isoforms in the included and excluded group, respectively, *I* denotes the set of all isoforms in gene *g*,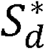 denotes the set of informative base pair positions of isoform *d*, and *S*_*d*_ denotes the set of all base pair positions of isoform *d*. To calculate weighted lengths of the region, we utilize observed read coverage *w*_*s*_ at base pair position *s* and relative abundance *θ*_*d*_ of isoform *d* which is estimated based on informative reads. Therefore, the numerator represents the weighted length of informative region for included exon group (+) and excluded exon group (-), respectively, and the denominator represents the weighted length of whole region in gene *g*.

### Detection of differential alternative splicing exon group

Exon-inclusion level, which measures the relative usage of an exon, is a commonly used measure to quantify the alternative splicing process. To detect DAS exons, we can examine if the inclusion level of exon group (*ψ*) is significantly different between two groups of cells. To make accurate inference on the inclusion level difference, technical parameters(*α*_*i*_, *β*_*i*_) for each cell *i* and population mean expression *θ*_*g*_ of gene *g* are first calculated during data pre-processing, using software TASC^18^. This allows us to eliminate bias due to technical variations in scRNA-seq data. Given an exon-group of interest, the informative read counts *Y*_+*i*_ /*Y*_−*i*_ are fitted to the hierarchical model described above, with likelihood function calculated as

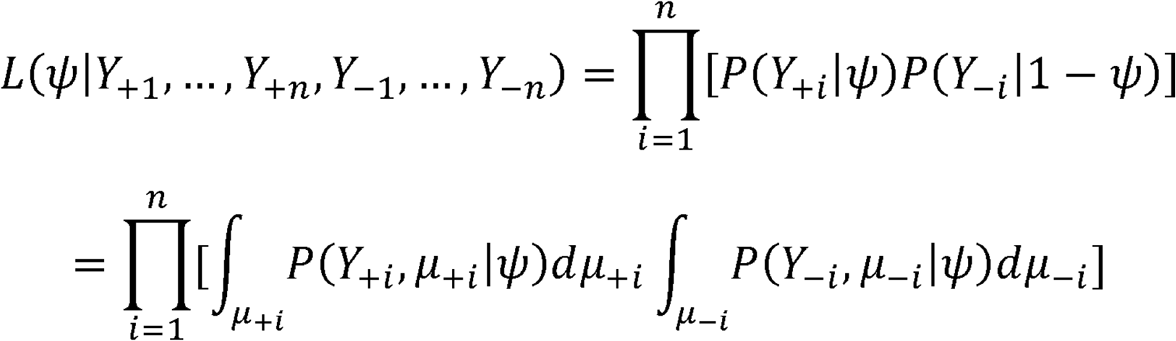

The joint distribution of cell-specific observed informative read counts and their true expression levels is

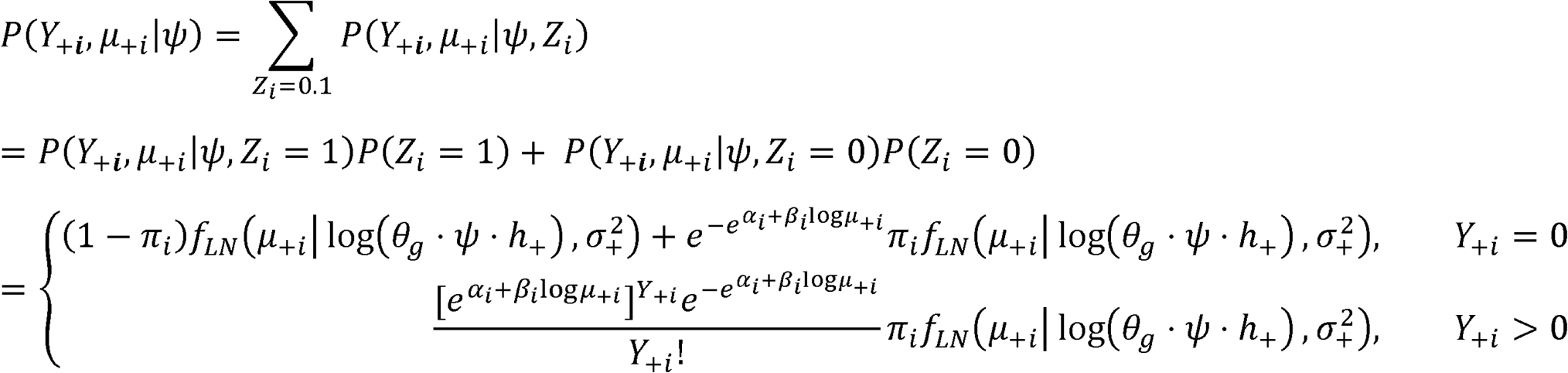

and similarly

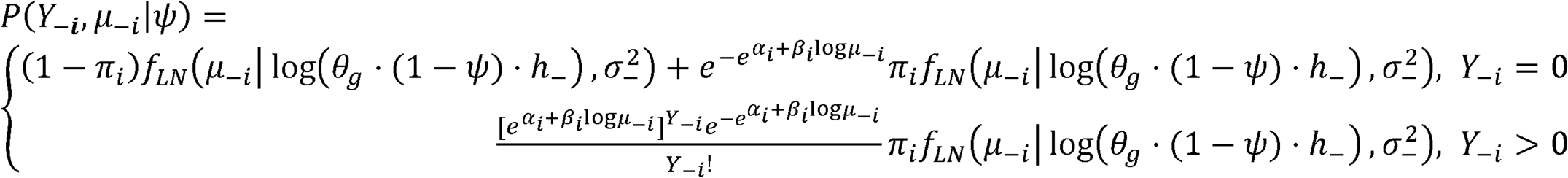

where the probability distribution function of log-normal distribution 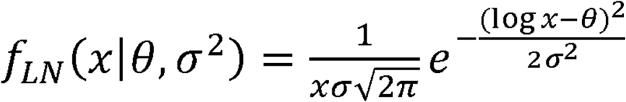.

To detect exons with DAS, we test for whether there is significant difference in inclusion level *ψ* between two group of cells, denoted by A and B, i.e.,*H*_0_: *ψ*_*A*_ = *ψ*_*B*_ vs. *H*_1_: *ψ*_*A*_ ≠ *ψ*_*B*_. A likelihood ratio test with test statistics 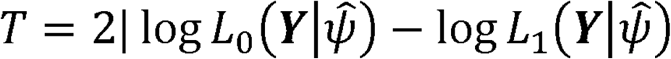 is employed to test this hypothesis, where 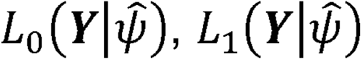 are maximized under the null hypothesis *H*_0_ and alternative hypothesis *H*_1_ respectively. Asymptotically, *T* follows a chi-square distribution with one degree of freedom.

### Datasets and evaluations

We first conducted a simulation study to benchmark the performance of SCATS and compared it with other existing DAS methods, including Census^10^ and DEXSeq^13^. To simulate realistic datasets, the average gene expressions (*θ*_*g*_) for each cell group were assigned based on measurements of human adipose bulk RNA-seq data^21^. The exon start and end positions were based on Ensembl annotation and downloaded in GTF format from UCSC Genome Browser (https://genome.ucsc.edu/). The cell-specific technical parameters (*α, β, κ, τ*) were estimated based on measurements from CA1 pyramidal cells in scRNA-seq data generated by Zeisel *et al*. 17. Given these abundances and technical parameters, we then parameterized the generative model described previously, which allows us to simulate exon-level informative read count (*Y*_+*ij*,_ *Y*_−*ij*_) mimicking the scRNA-seq data generation process. Following this procedure, we simulated 1,000 expressed genes with 6,000 total reads per cell for 600 cells (300 in condition A, and 300 in condition B) for five scenarios (Δ = | *ψ*_*A*_ − *ψ*_*B*_ | = 0, 0.1, 0.2, 0.3, 0.4) to evaluate type I error rate and power for SCATS, Census and DEXSeq.

Next, we evaluated different DAS detection methods using mouse cortical dataset generated by Tasic *et al*.^14^, which includes 1,679 cells from 49 cell types in three major cell classes (23 GABAergic cell types, 19 glutamatergic cell types, and seven non-neuronal cell types). Among these cells, the average sequencing depth is 3 million. We directly downloaded the aligned data in BAM format from NCBI Gene Expression Omnibus (<u>GSE71585</u>). Given read counts of 6,275 endogenous RefSeq genes with at least two annotated isoforms and 92 ERCC spiking-ins across cells, we applied TASC to quantify technical variations, yielding accurate estimations of average gene concentrations in each sub cell type. Similarly, informative reads in each exon group were counted using SCATS based on the 6,275 genes with 12,073 exon groups. On average, 4,311 exon groups were covered by at least one splicing informative read in each cell. We treat an exon group as qualified for analysis when at least three cells have at least one informative read in each condition. Pairwise DAS comparison was conducted between these 49 sub cell types with 985 qualified exon group on average for each comparison.

Another adult mouse brain data used for evaluation was acquired from Zeisel *et al*.^17^, which includes sequencing data of 3,005 cells from various regions of mouse brain. These cells were classified into nine major cell classes and 47 sub cell types, with the sub cell types from the same major cell type considered to be relatively homogenous. In this study, raw data in FASTQ format were downloaded from NCBI Gene Expression Omnibus (<u>GSE60361</u>) and aligned to mm10 reference genome with Ensembl gene annotation using STAR^22^. For each cell, we processed the BAM file and counted splicing informative reads using SCATS based on 1,826 marker genes of nine major cell types (with at least two annotated isoforms) with 3,542 exon groups annotated by Ensembl. For each cell, there are 465 exon groups on average covered by at least one informative UMI read. Given these informative counts, DAS analysis between each pair of these 48 sub cell types was conducted. To compare the ability of DAS detection, similarly, we performed pairwise DAS analysis across sub cell types using MISO^16^. For each comparison, we required at least three cells have at least one informative UMI in each condition, leading to 568 qualified exon groups on average for each DAS comparison.

## Software availability

An open-source implementation of the SCATS algorithm can be downloaded from https://github.com/huyustats/SCATS.

## Life sciences reporting summary

Further information on experimental design is available in the Life Sciences Reporting Summary.

## Data availability

The published data sets used in this manuscript are available through the following websites or accession numbers: GSE60361 (Zeisel *et al*. data) and GSE71585 (Tasic *et al*. data).

