## Supplementary materials for "Detecting differential alternative splicing events in scRNA-seq with or without UMIs"

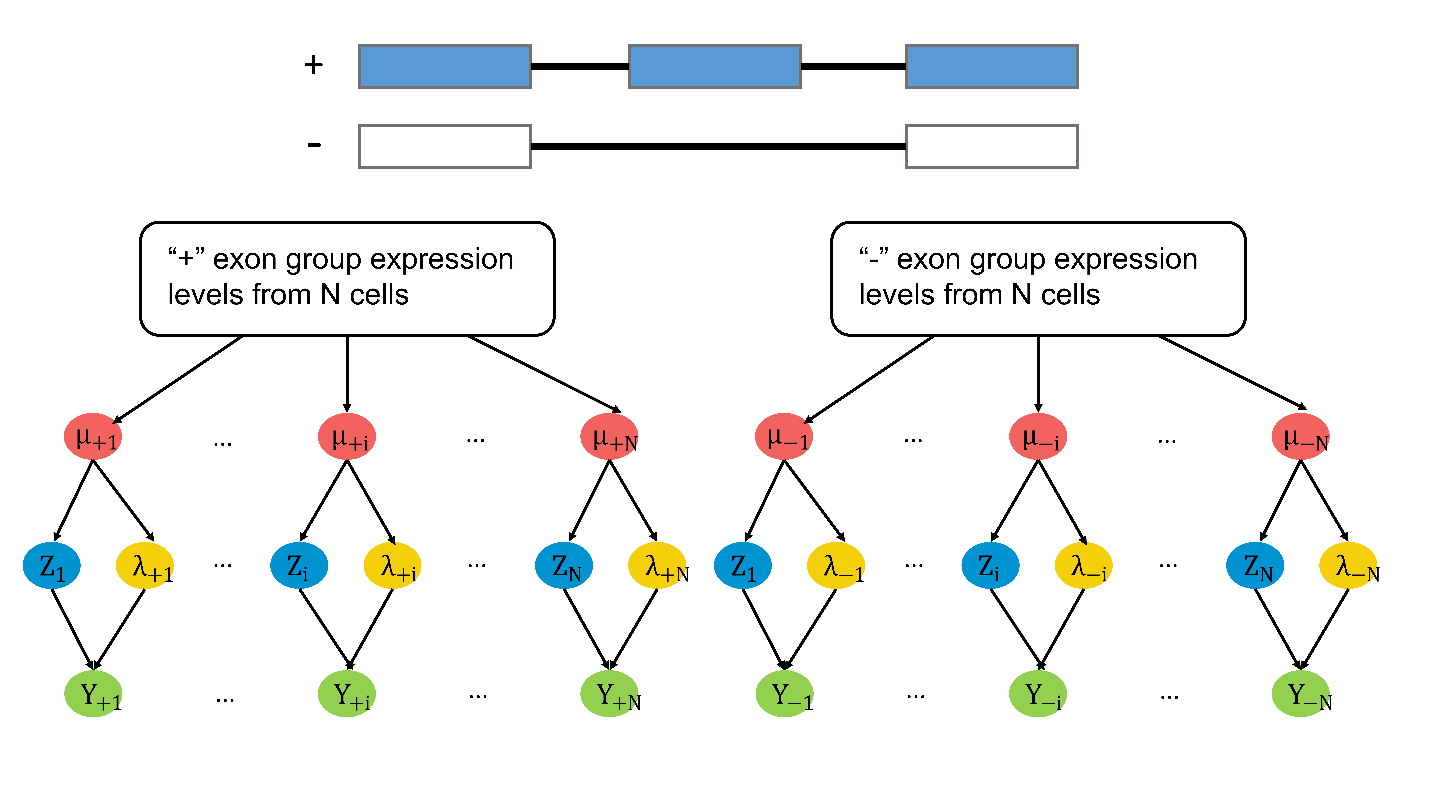

**Supplementary Fig. 1: Schematic of the SCATS model.** An alternatively spliced exon is included in the “+” exon group (blue) and excluded from the “-” group (white). $\mu_{+i}$ and $\mu_{-i}$ represent true expression level of the “+” and “-” exon groups in cell $i$, respectively. $\lambda_{+i}$ and $\lambda_{-i}$ are intermediate variables that model amplification bias, capture efficiency, and sequencing bias in cell $i$ for the “+” and “-“ exon groups. $Z_{i}$ models transcriptional bursting and dropout event of the gene in cell $i$. $Y_{+i}$ and $Y_{-i}$ represent observed informative read counts of the “+” and “-” exon groups in cell $i$. SCATS utilizes a hierarchical model to detect differential alternative splicing (DAS) event between cell groups by accounting for technical noise from single-cell RNA-seq data.

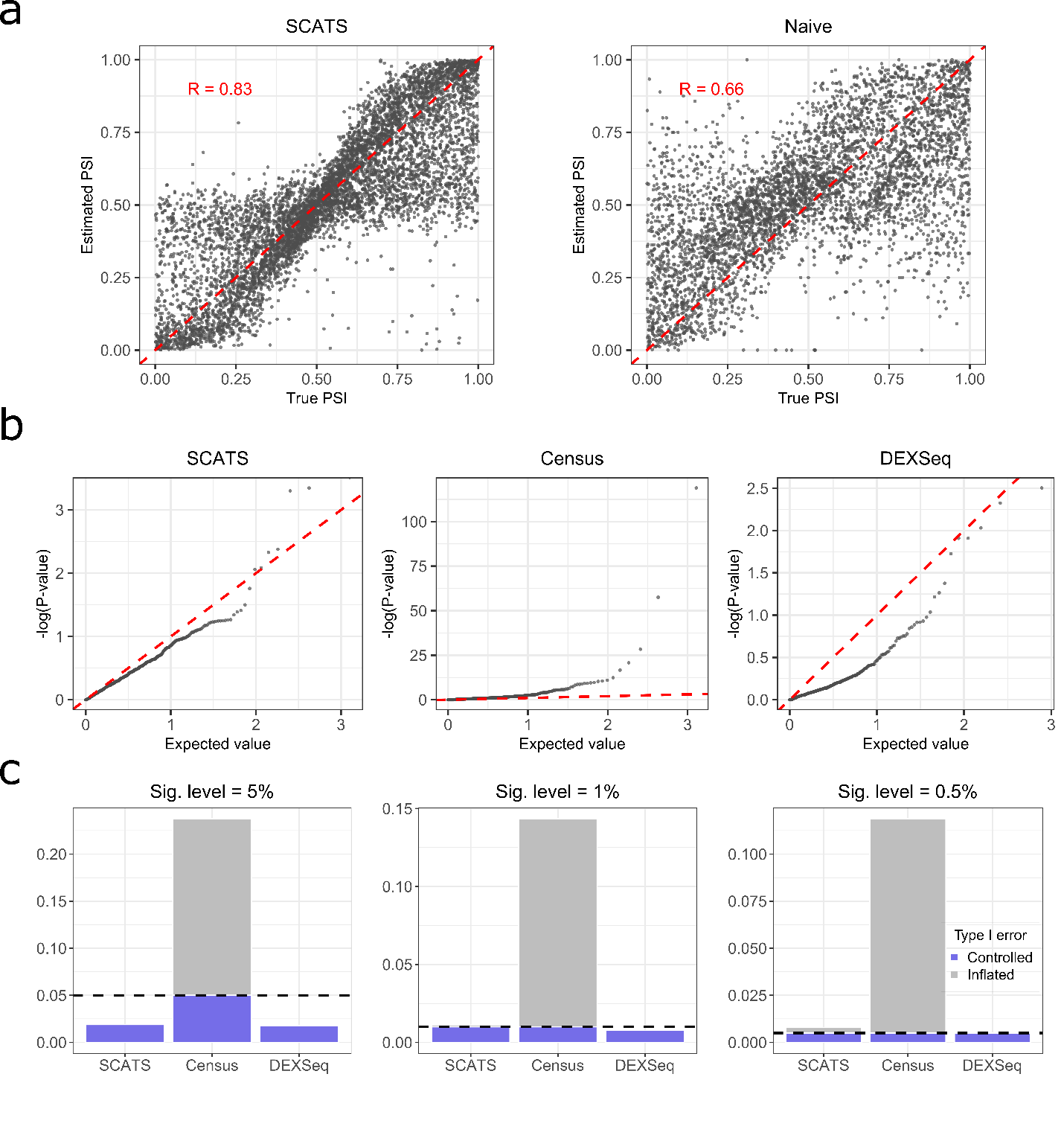

**Supplementary Fig. 2: Exon-inclusion level estimation and false positive rate comparison.** Informative read counts of 1,000 exon groups for 600 cells (300 vs 300) were simulated using generative model shown in **Supplementary Fig. 1** with cell-specific technical parameters estimated from CA1 pyramidal cells in the scRNA-seq data generated by Zeisel *et al*.^1^. To simulate data at resemble real scRNA-seq data, one CA1 pyramidal cell was randomly selected and read coverage for each gene in that cell was used to obtain true gene concentration at the population level ($gene concentration=100,000\times\frac{\# reads from the gene}{\# total number of reads from the CA1 pyramidal cell}$). Based on the generative model accounting for technical noise, we finally generated 6,000 reads on average for each cell. **(a)** Scatter plots of 1,842 true exon-inclusion levels, estimated by percent spliced in (PSI) for a given exon, against estimates from SCATS and naïve method that ignores technical noise. SCATS models technical noise to quantify usage of an alternatively spliced exon while the naïve method simply estimates the inclusion level by $\psi=\frac{Y_{+}}{Y_{+}+Y_{-}}$, where $Y_{+}$ and $Y_{-}$ are the informative read counts of the “+” and “-” exon groups across cells. SCATS estimates are closer to the ground truths. **(b)** Quantile-quantile plots of the p-values from SCATS, Census and DEXSeq under the null hypothesis ($\Delta=0$). X-axis represents uniform theoretical quantiles between 0 and 1 in ${-\log}_{10}$scale. Y-axis represents observed p-value quantile in ${-\log}_{10}$scale. Uniformly distributed data should follow the red dashed line. P-values of SCATS are more uniformly distributed while Census being right-skewed and DEXSeq being left-skewed. **(c)** Type I error comparison of SCATS, Census, and DEXSeq with different significance levels (α = 0.05, 0.01, 0.005). Consistent with **(b)**, SCATS has better type I error control than Census and DEXSeq.

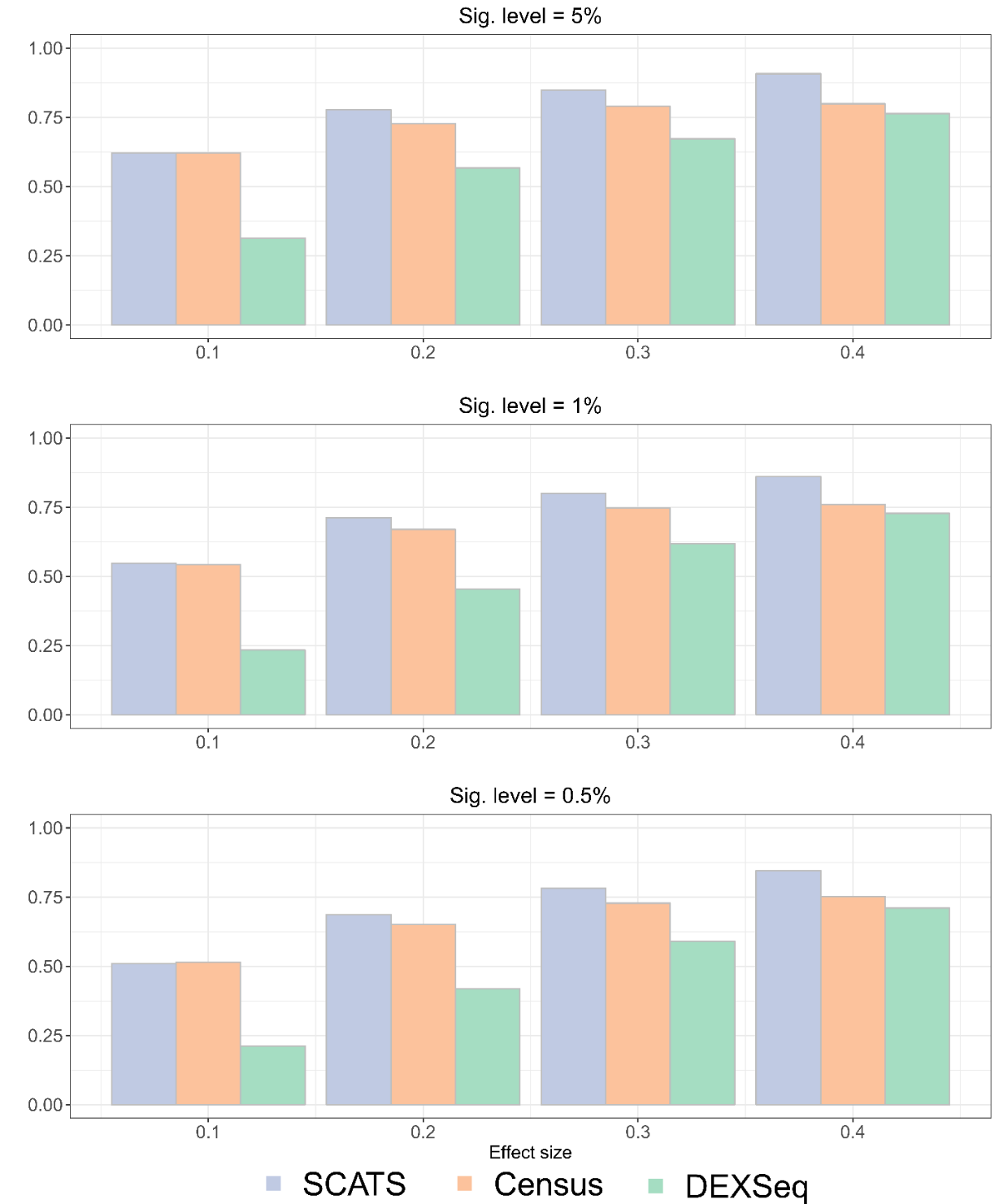

**Supplementary Fig. 3: Power comparison of SCATS, Census and DEXSeq.** Informative read counts of 1,000 exon groups for 600 cells (300 vs 300) were simulated using generative model shown in **Supplementary Fig. 1** with cell-specific technical parameters estimated from CA1 pyramidal cells in the scRNA-seq data generated by Zeisel *et al*.^1^. The average sequencing depth per cell is 6,000. Barplots show the estimated power under different effect sizes ($\Delta=0.1, 0.2, 0.3, 0.4$). Significance was evaluated at 0.05, 0.01, and 0.005 levels. Colors indicate different methods. SCATS outperforms Census and DEXSeq across all effect sizes, especially when $\Delta=0.1$. DEXSeq is conservative in detecting DAS events.

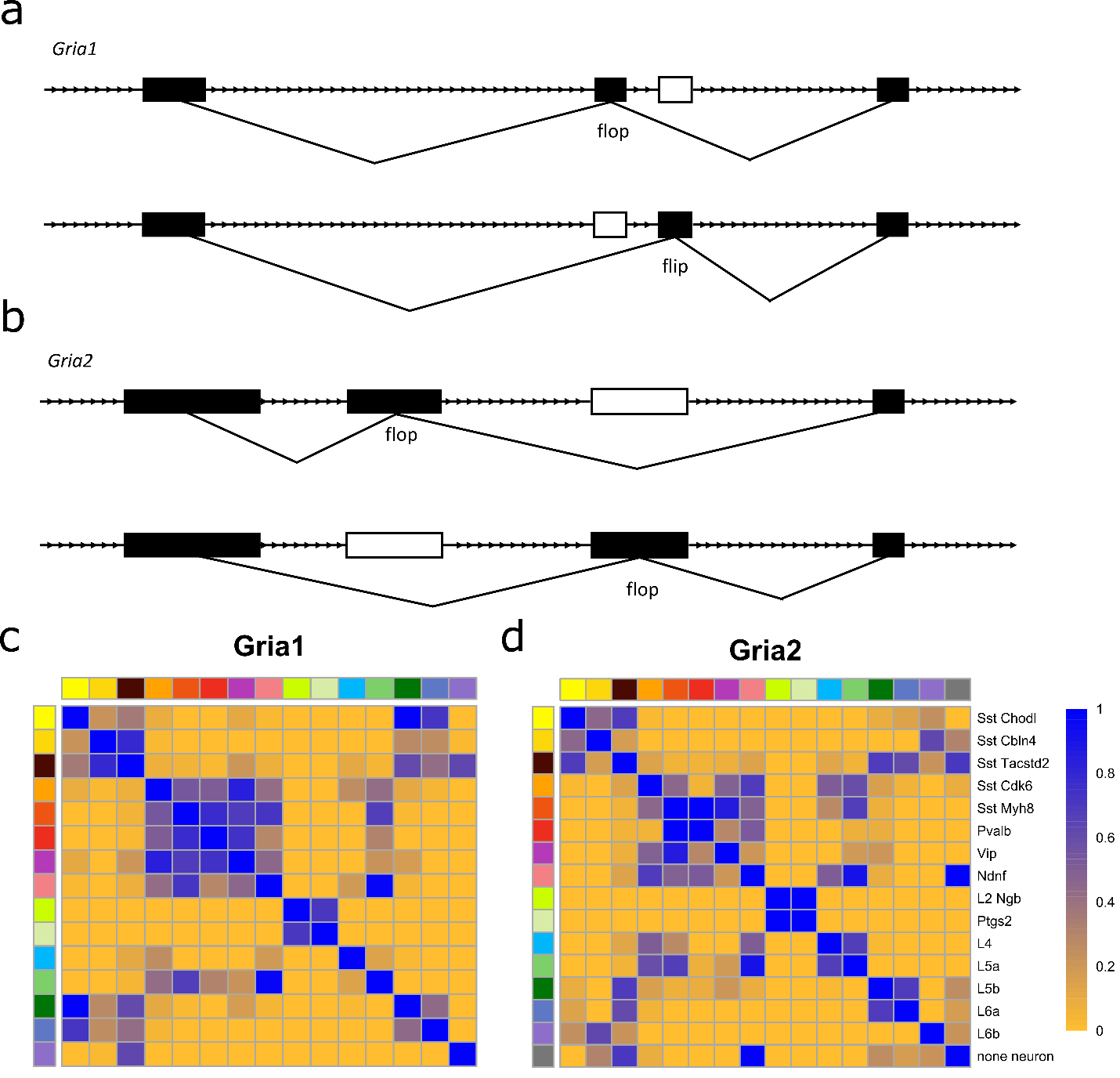

**Supplementary Fig. 4: DAS analysis of AMPA receptor genes, *Gria1* and *Gria2* using Tasic *et al.* dataset. (a,b)** Mutually exclusively alternatively spliced flip-flop exons of genes *Gria1* **(a)** and *Gria2* **(b)**. **(c,d)** Heatmap showing the p-values of pairwise DAS tests on the flop exon for gene *Gria1* **(c)** and *Gria2* **(d)** across 16 cell types (8 GABAergic, 7 Glutamatergic, 1 non-neuronal) using SCATS. These 16 cell types were selected by Tasic et al. in which they found highly cell type-specific splicing patterns across these cell types. As expected, SCATS results also showed highly cell type-specific splicing pattern of the *Gria* genes.

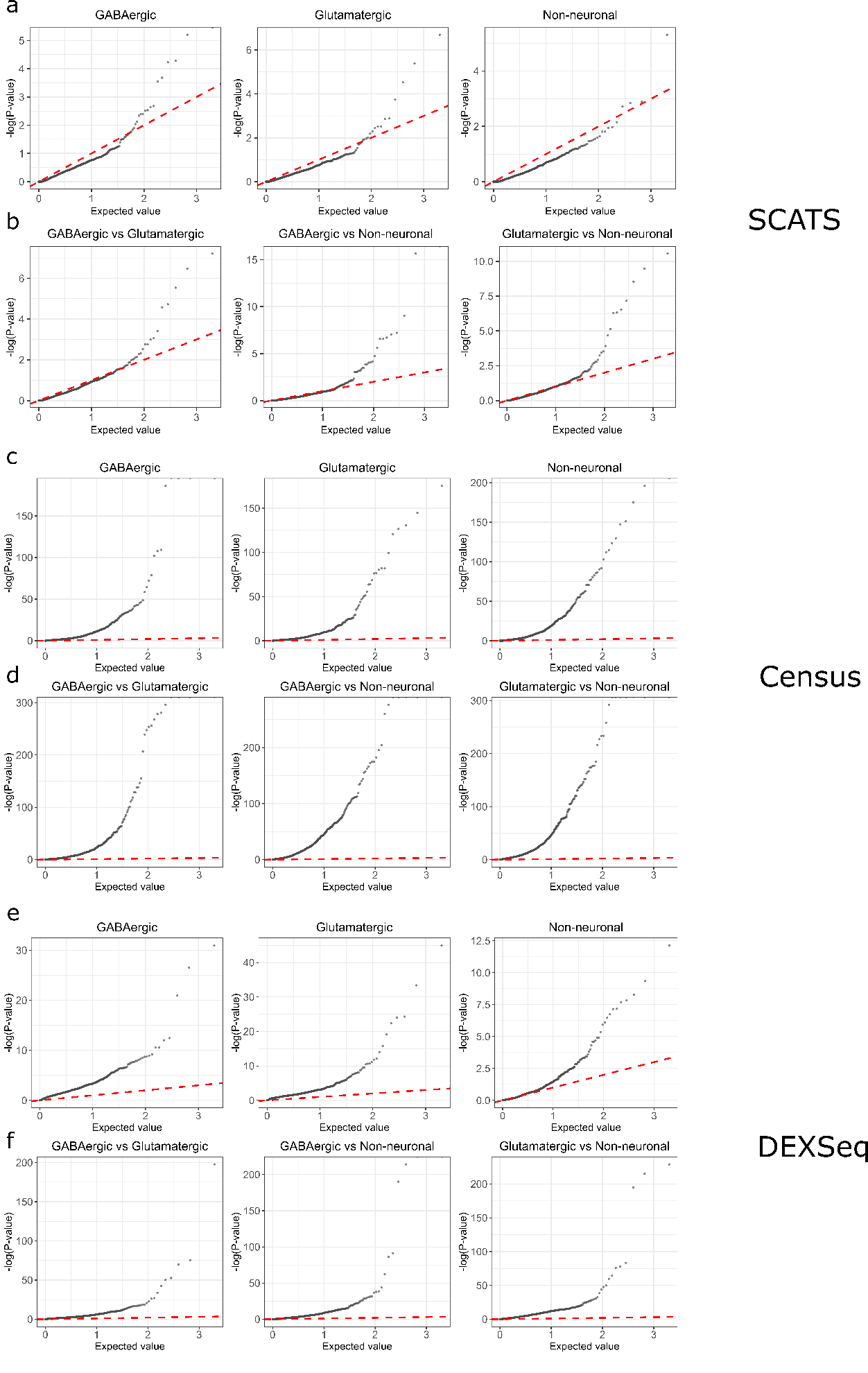

**Supplementary Fig. 5: Quantile-quantile plots of p-values from SCATS, DEXSeq and Census on the Tasic *et al*. dataset^2^.** Pairwise DAS comparison across 49 cell types in three major cell classes was performed using SCATS. DAS comparisons were classified into two groups: comparison for cell types within major cell classes **(a, c, e)**, and comparison for cell types across major cell classes **(b, d, f)**. X-axis represents uniform theoretical quantiles between 0 and 1 in ${-\log}_{10}$scale. Y-axis represents observed p-value quantile in ${-\log}_{10}$scale. Uniformly distributed data should follow the red dashed line. QQ plots of p-values from within major cell class comparisons (GABAergic, Glutamatergic, Non-neuronal) are similar to the plot under the null hypothesis in simulation study. This matched our expectation because most exons should not be differentially spliced between cell types within the same major cell class. In contrast, distribution of p-values is highly right-skewed starting from X=10^-1.5^ for cross-major cell class comparisons (GABAergic vs. Glutamatergic, GABAergic vs. Non-neuronal, Glutamatergic vs. Non-neuronal), indicating that more DAS events were detected.

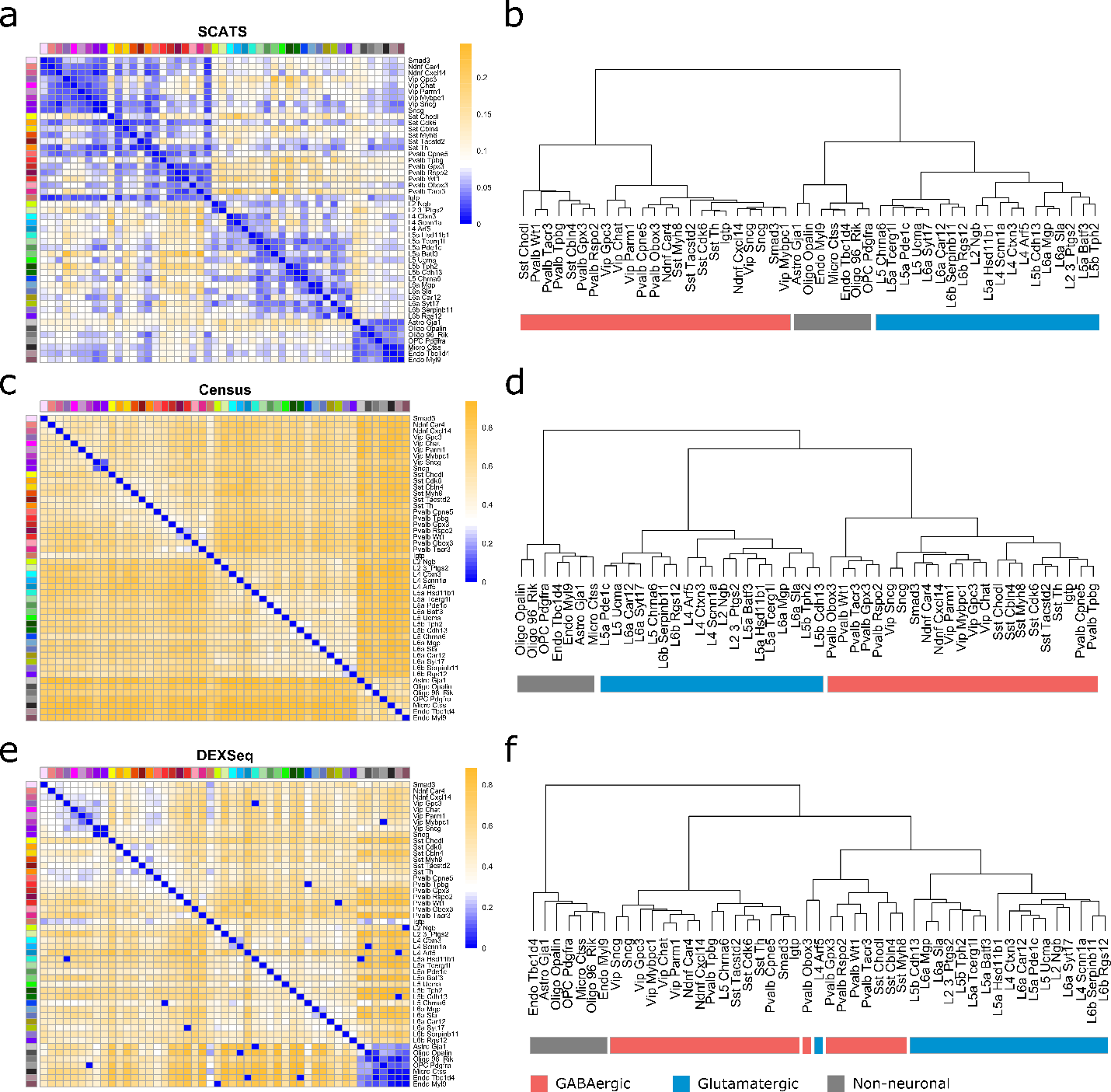

**Supplementary Fig. 6: DAS analysis results of the Tasic *et al*. data^2^ based on 296 genes with 966 exon groups.** The 296 genes were detected as the most differentially spliced genes by Tasic et al. based on MISO analysis. **(a-f)** Pairwise DAS comparison across 49 cell types from three major cell classes: GABAergic, Glutamatergic, Non-neuronal from mouse cortex using SCATS **(a,b)**, Census **(c,d)** and DEXSeq **(e,f)**. Colors indicate different cell types. **(a,c,e)** Heatmap showing the proportion of detected DAS exon groups for each pairwise comparison between cell types. **(b,d,f)** Dendrogram depicting cell classification results of the 49 cell types. Each cell type was marked by the corresponding major cell class: GABAergic (red), Glutamatergic (blue) and Non-neuronal (grey). The distance metric between two cell types is the proportion of detected DAS exon groups among all analyzed exon groups by each method: SCATS **(b)**, Census **(d)** or DEXSeq **(f)**. SCATS and Census yield better cell type classification result than DEXSeq.

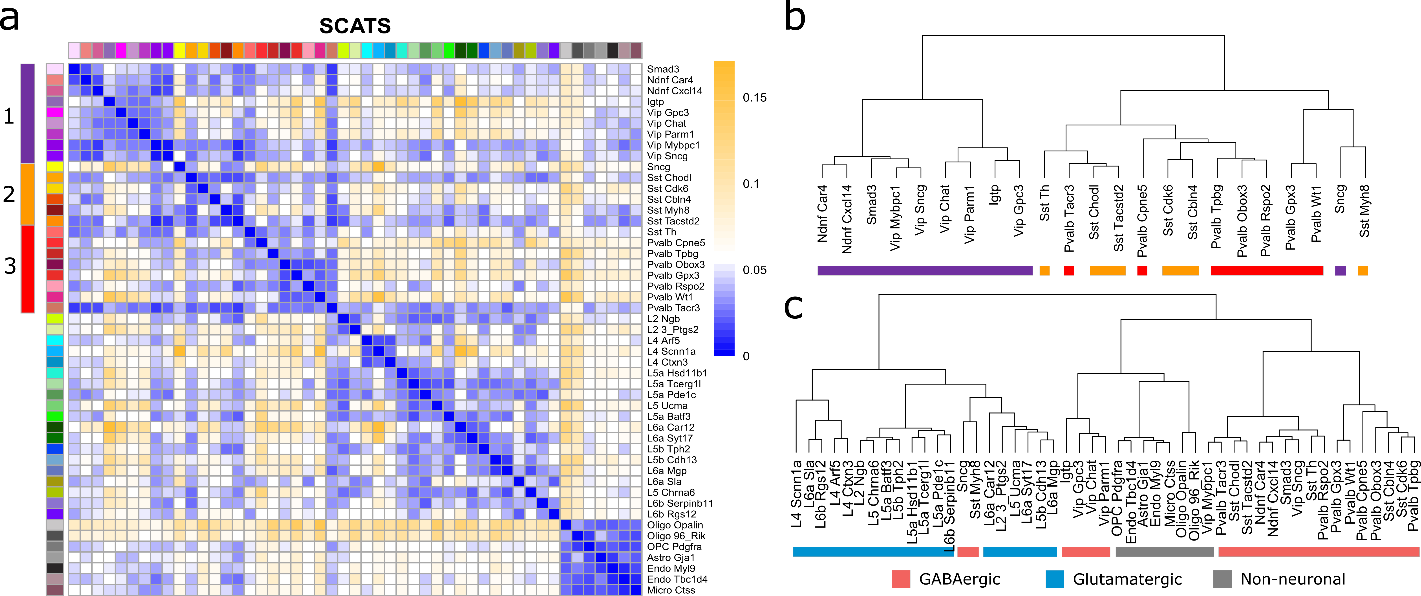

**Supplementary Fig. 7: DAS analysis results of the Tasic *et al*. data^2^ based on 6,275 genes with 12,073 exon groups using SCATS.** Pairwise DAS comparison across 49 cell types from three major cell classes: GABAergic, Glutamatergic and Non-neuronal. **(a)** Heatmap showing the proportion of detected DAS exons groups for each pairwise comparison. **(b,c)** Dendrogram depicting cell classification results of the 23 GABAergic cell types **(b)** and all 49 cell types **(c)**. The distance metric between two cell types is the proportion of detected DAS exon groups among all analyzed exon groups by SCATS. **(b)** Each GABAergic cell type was marked by the corresponding cell type (red, yellow, purple). These cell types were validated based on hierarchical clustering analysis of qRT-PCR measurements of 79 marker genes by Tasic *et al*. [2]. **(c)** Each cell type was marked by the corresponding major cell class: GABAergic (red), Glutamatergic (blue) and Non-neuronal (grey).

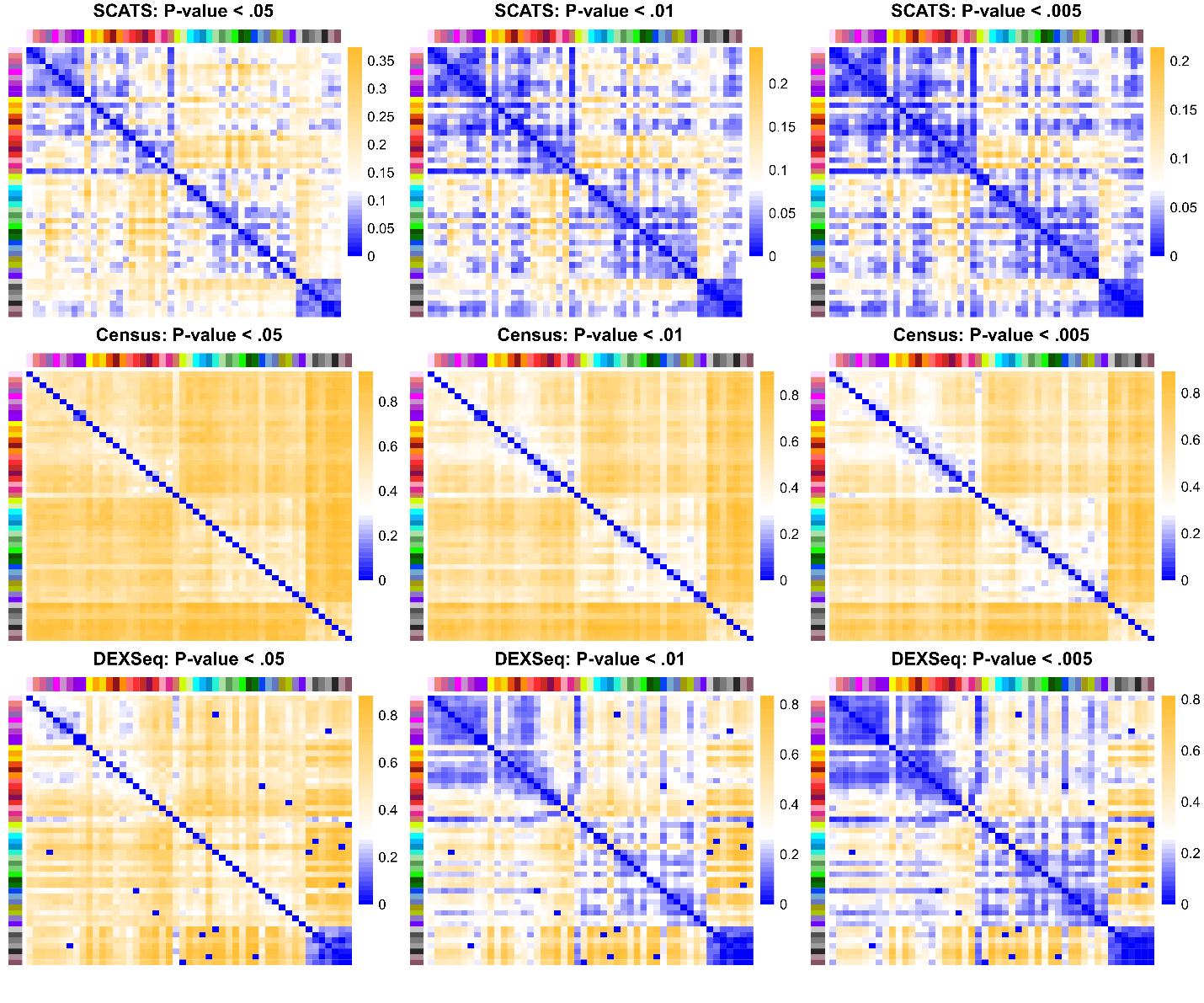

**Supplementary Fig. 8: DAS analysis comparison across different significance levels and methods.** The analysis was conducted on 296 genes detected as the most differentially spliced genes by Tasic et al. based on MISO analysis. Heatmap showing the proportion of detected DAS exons groups for each pairwise comparison between 49 cell types. DAS detection performance of SCATS, Census and DEXSeq was evaluated using different significance levels (α = 0.05, 0.01, 0.005). For within major cell class comparisons (GABAergic, Glutamatergic, Non-neuronal), Census yielded much higher DAS detection (~0.6) rate than SCATS (<0.1), indicating possible inflated false positive rate for Census.

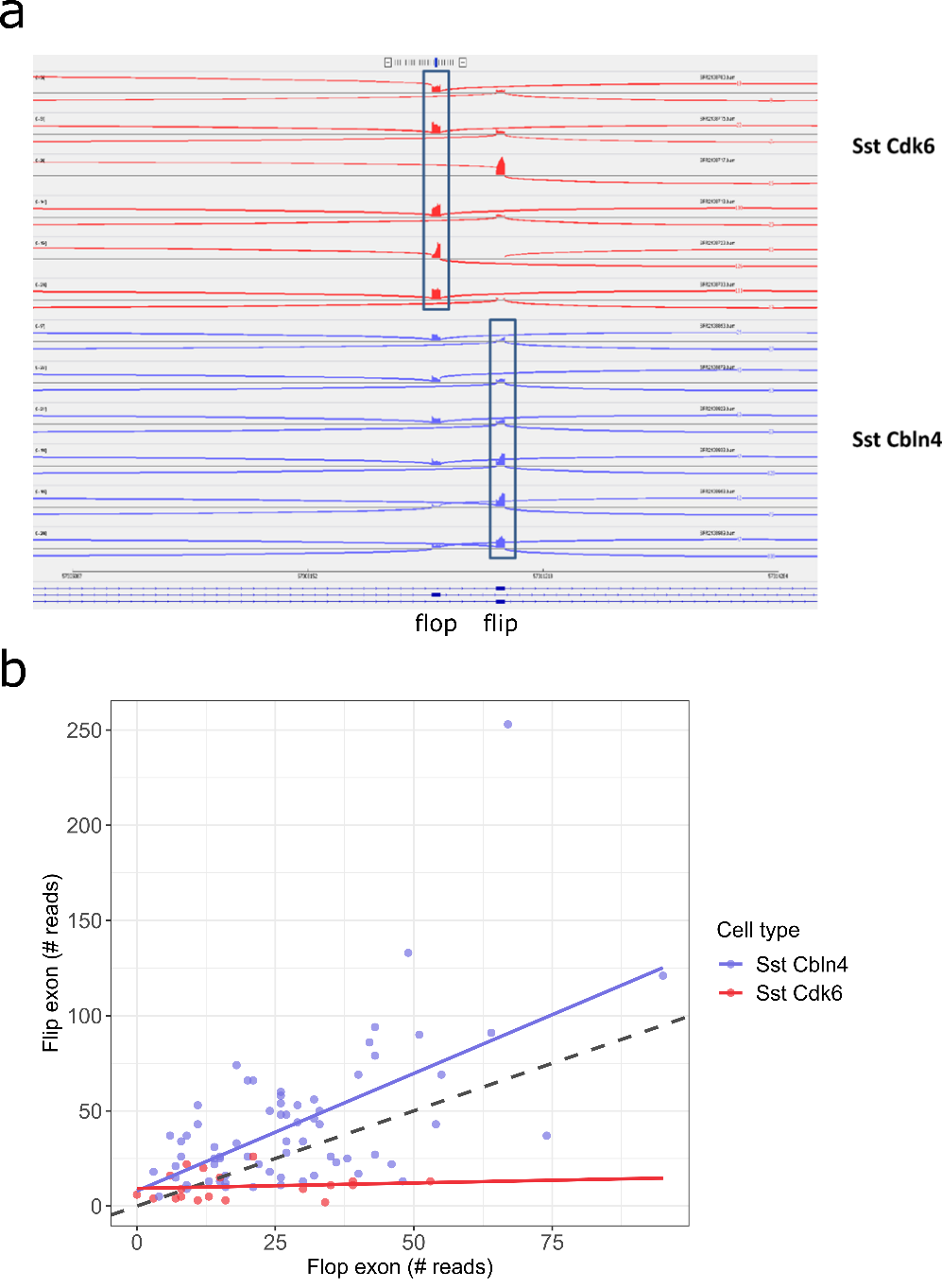

**Supplementary Fig. 9: Informative read coverage of flip-flop exon of gene *Gria1* based on Tasic *et al.* data. (a)** IGV sashimi plot of flip-flop exon between cell types Sst Cdk6 and Sst Cbln4. This differentially spliced flip-flop was identified by SCATS but missed by Census and DEXSeq. Each row represents one single cell. For each cell type, 6 cells were randomly selected respectively (68 Sst Cbln4 cells, 19 Sst Cdk6 cells). **(b)** Scatter plot of flip exon coverage against flop exon coverage across all 87 cells. Black dashed line is identical line. Read coverages showed a significant difference in flip-flop exon usage between cell types Sst Cdk6 and Sst Cbln4.

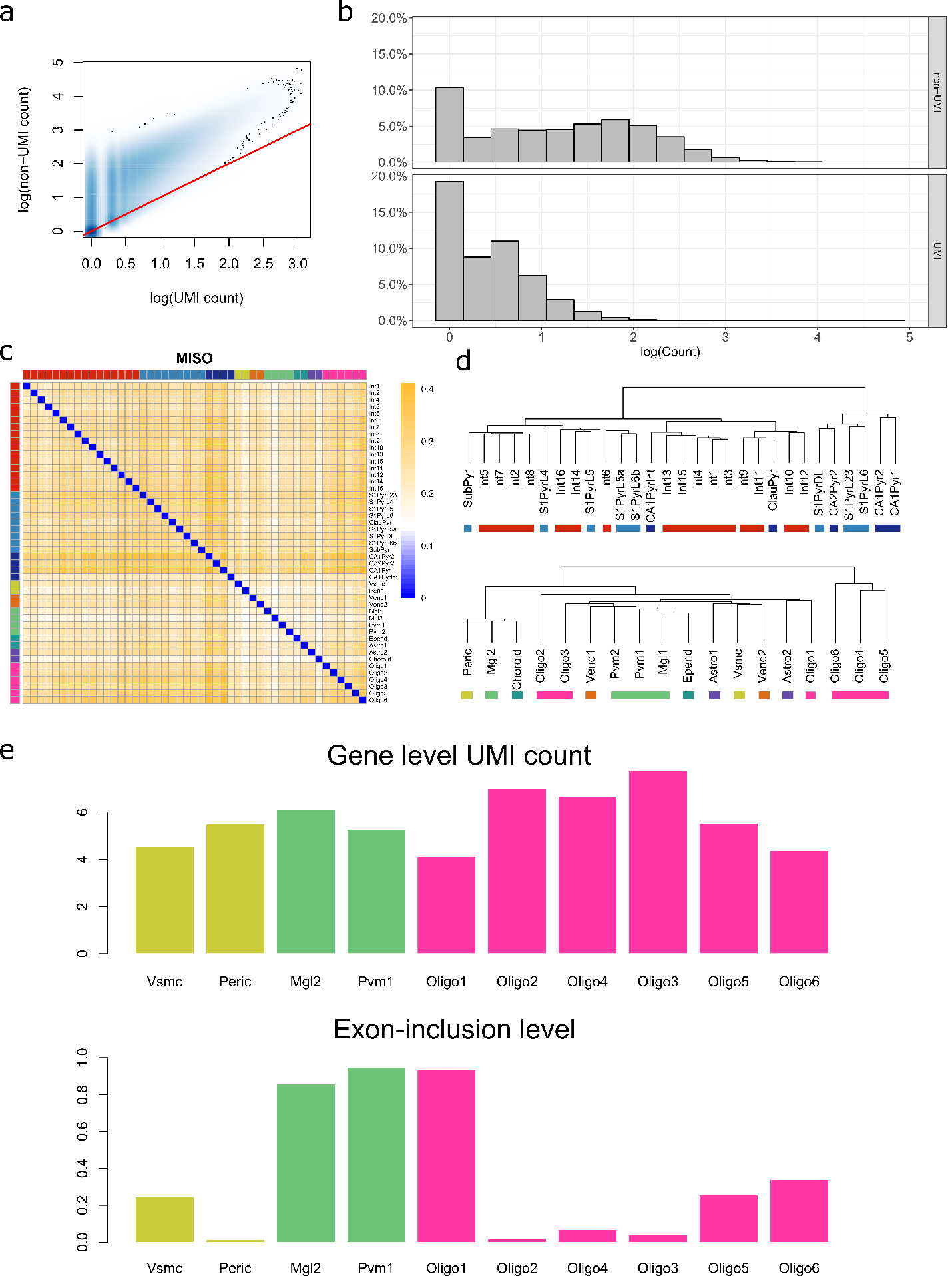

**Supplementary Fig. 10: DAS analysis results of the Zeisel *et al*. data.** This dataset includes both UMI and non-UMI read counts. The UMI read counts were obtained by consolidating read counts for reads that originate from the same molecule based on UMI barcodes. This allows us to compare UMI and non-UMI read counts systematically for DAS analysis. **(a,b)** Comparison of UMI and non-UMI read counts. **(a)** Scatter plot of UMI splicing informative counts against non-UMI splicing informative counts in log scale. Each point was summarized based on the same exon group in the same cell. Red dashed line represents identical line. **(b)** Distribution comparison between UMI and non-UMI splicing informative read counts. UMI counts are much sparser than non-UMI counts, suggesting that splicing analysis using UMI counts is more challenging. **(c,d)** Pairwise DAS comparison across nine major cell types from mouse cortex and hippocampus using MISO. Colors indicate nine major cell types. **(c)** Heatmap showing the proportion of detected DAS exon groups for each pairwise comparison between cell types. **(d)** Dendrogram depicting cell classification results of the 47 sub cell types. The distance metric between two sub cell types is the proportion of detected DAS exons among all analyzed exons by MISO. MISO showed the worst performance in cell type classification as compared to SCATS, Census and DEXSeq in **Fig. 3**. **(e)** Gene-level UMI counts and exon-inclusion level estimates of the flip exon from gene *Gria1* across cell types. Splicing quantification offers higher resolution of cellular heterogeneity than regular gene expression.

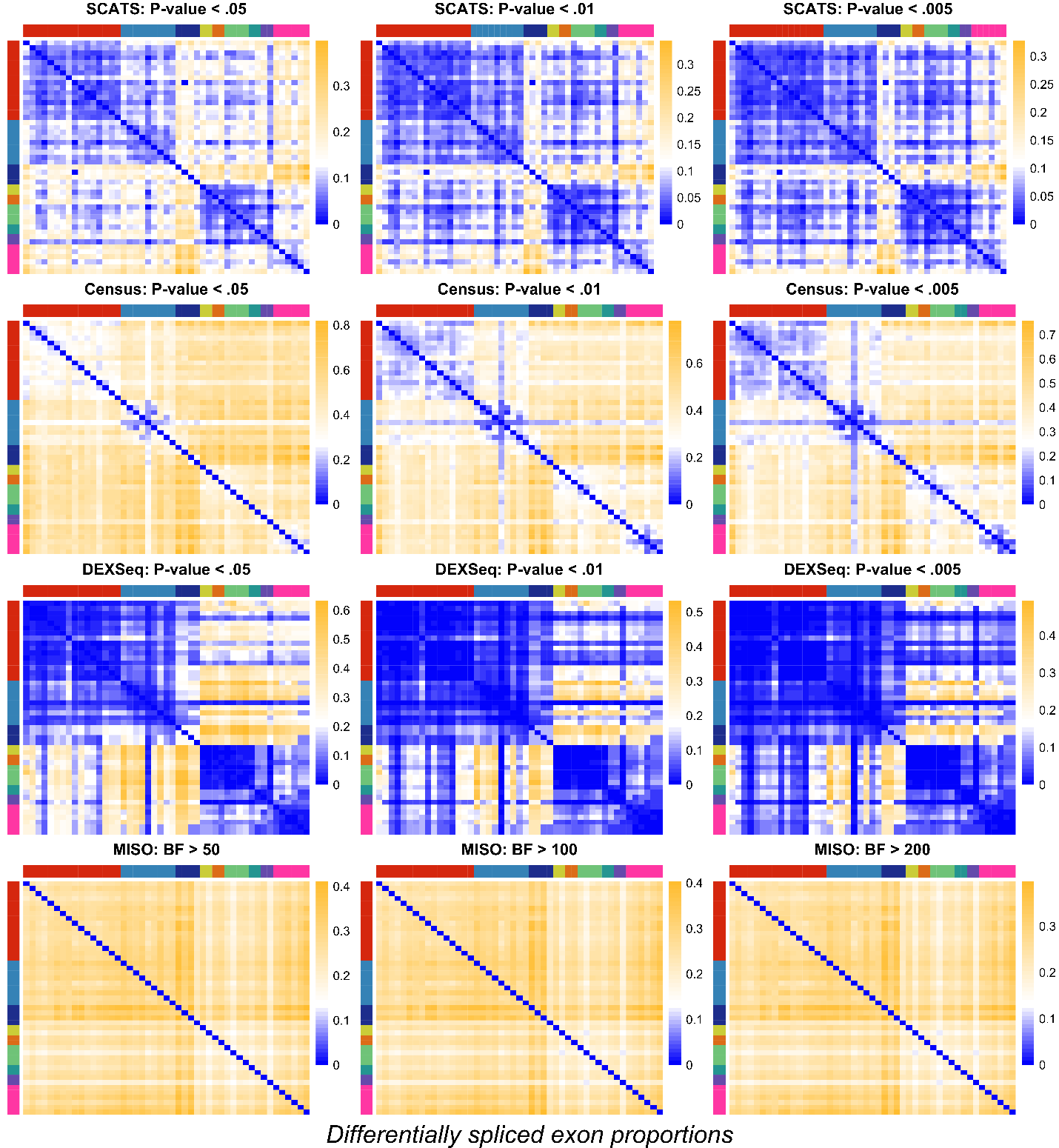

**Supplementary Fig. 11: DAS analysis comparison across different significance levels and methods using Zeisel *et al.* data.** Heatmap showing the proportion of detected DAS exon groups (3,542 exon groups from 1,826 genes) for each pairwise comparison between nine major cell types. DAS detection performance of SCATS, Census, DEXSeq and MISO was evaluated using different significance levels (p-value α=0.05, 0.01, 0.005 or Bayes factor: 50, 100, 200) respectively.
